# ATP emerged to induce protein folding, inhibit aggregation and increase stability

**DOI:** 10.1101/739581

**Authors:** Jian Kang, Liangzhong Lim, Jianxing Song

**Affiliations:** Department of Biological Sciences, Faculty of Science, National University of Singapore; 10 Kent Ridge Crescent, Singapore 119260

**Keywords:** ATP, Triphosphate, Protein folding, Protein aggregation, Protein dynamics, Amyotrophic lateral sclerosis (ALS), NMR spectroscopy

## Abstract

By NMR characterization of effects of ATP and related molecules on the folding and dynamics of the ALS-causing C71G-PFN1 and nascent hSOD1, we reveal for the first time that ATP has a general capacity in inducing protein folding with the highest efficiency known so far. This capacity was further identified to result from triphosphate, a key intermediate in prebiotic chemistry, which, however, can severely trigger protein aggregation. Remarkably, by joining adenosine and triphosphate together, ATP integrates three abilities to simultaneously induce protein folding, inhibit aggregation and increase thermodynamic stability. Our study implies that the emergence of ATP might represent an irreplaceable step essential for the Origin of Life, and decrypts a principle for engineering small molecules with three functions to treat aggregation-associated ageing and diseases.

**One sentence summary:** By joining adenosine and triphosphate, ATP integrates three abilities to control protein homeostasis for the Origin of Life.

---

Cell is extremely crowded in which protein concentrations can exceed 100 mg/ml, ~3 mM of an ‘average’ protein (1). Many proteins also need to fold from the unfolded state (U) into the native state (N), although some remain intrinsically disordered (2–10). Protein aggregation is the hallmark of aging and neurodegenerative diseases including amyotrophic lateral sclerosis (ALS) (10–14). Recently it was decoded that ATP, the universal energy currency for all cells with very high concentrations (1-12 mM) (15), can act as a hydrotrope to prevent/dissolve protein aggregation (12,13). We additionally found that by specific binding, ATP modulates liquid-liquid phase separation (LLPS) (16,17), inhibits fibrillation (18), and antagonizes pathological aggregation (19). However, all these functions require very high ATP concentrations and therefore one argument is that even the maximal ATP cellular concentration (~12 mM) only accounts for ~4 ATP/protein, not to mention that in neurons, ATP concentration is only ~3 mM (12,13,15).

Here we thus asked whether ATP also evolved to modulate protein folding process. As the native state is only marginally stable, even in the modern cells with the ATP-energy-dependent chaperone machineries, genetic mutations or even absence of co-factors are sufficient to render the native protein to co-exist with the unfolded state or to become completely unfolded, which consequently become prone to aggregation mainly due to the electrostatic screening effect in salted buffers (10–14). To address this question, we first studied the effect of ATP on the folding equilibrium of the ALS-causing C71G mutant of profilin-1 (PFN1). PFN1 is an 140-residue protein adopting a seven stranded antiparallel β-sheet sandwiched by α-helices (Fig. 1A). C71G-PFN1 characteristic of severe aggregation causes ALS by gain of toxicity (20). Previously we showed that C71G-PFN1 co-existed in equilibrium between the unfolded and folded states (21), as characterized by two set of HSQC peaks (Fig. S1A). Here we successfully achieved NMR assignments of WT-PFN1 and C71G-PFN1 except for the missing or overlapping peaks. WT-PFN1 and the folded state of C71G-PFN1 have highly similar (ΔCα-ΔCβ) values (Fig. S2A), indicating that they have very similar conformations. However, the absolute values of (ΔCα-ΔCβ) of the unfolded state are much smaller than those of its folded state (Fig. S2B), clearly suggesting that the unfolded state is highly disordered without any stable secondary and tertiary structures.

**Fig. 1.**
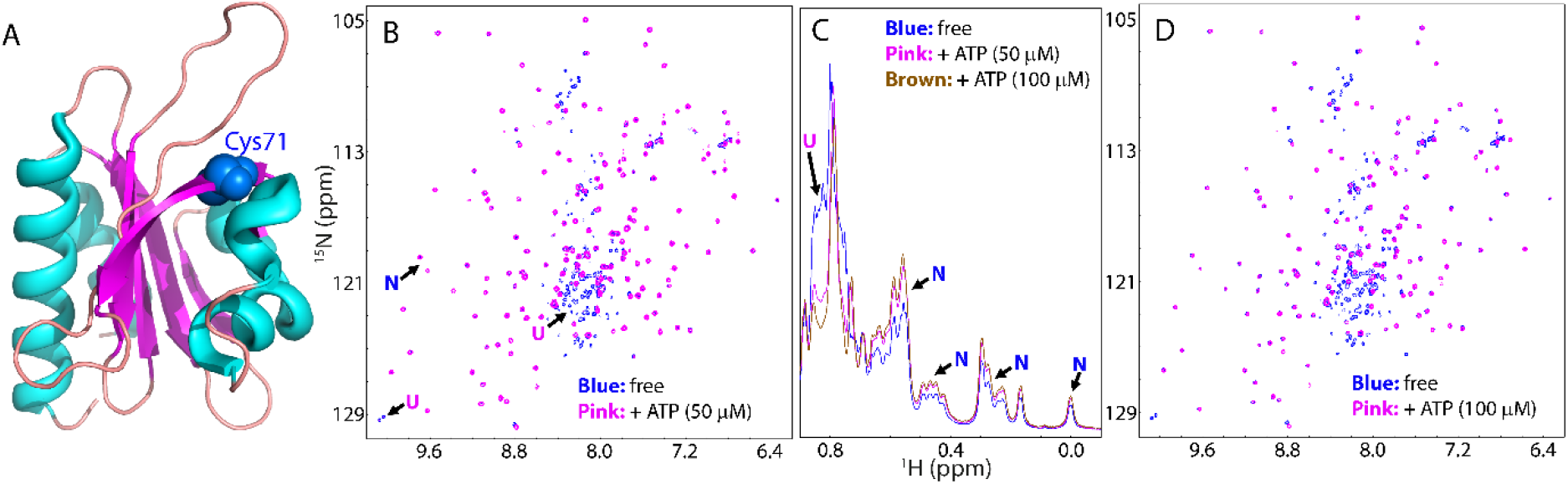
ATP shifts the conformational equilibrium to favor the folded state. (A) The three-dimensional structure of the human profilin-1 (PDB ID of 2PAV) with the Cys71 residue displayed in blue spheres. (B) Superimposition of HSQC spectra of ^15^N-labeled C71G-PFN1 at a concentration of 50 μM in the absence (blue) or ATP (pink) at a molar ratio of 1:1. (C) Up-field 1D NMR spectra of C71G-PFN1 in the presence of ATP at different ratios. (D) Superimposition of HSQC spectra of C71G-PFN1 in the absence (blue) and in the presence of ATP (pink) at a molar ratio of 1:2. Some characteristic NMR signals of the folded native (N) and unfolded (U) states were indicated by arrows.

Strikingly, when ATP was added even only at a molar ratio of 1:0.5 (C71G:ATP), the intensity of HSQC peaks of the unfolded state became reduced and those of the folded state slightly increased (Fig. S1B and S3A). The 1D peak intensity of the methyl group from the unfolded state also became reduced while those of the folded state slightly increased (Fig. S1C). When ATP was added to 1:1, the HSQC peak intensity of the unfolded state became further reduced while those of the folded state increased (Fig. 1B and S3B). At 1:2, HSQC peaks of the unfolded state became completely disappeared (Fig. 1D), and the 1D peak intensity of the methyl group of the folded state increased (Fig. 1C). Further addition of ATP to 1:20 (1 mM) showed no significant change of HSQC spectrum (Fig S1D). These results indicate that ATP has the very strong capacity in shifting the conformational equilibrium to favor the folded state. We also increased the C71G-PFN1 concentration to 100 μM, and the ATP concentration required to completely convert the unfolded state also doubled (200 μM). This suggests that the conversion by ATP is dependent of the molar ratio between C71G-PFN1 and ATP.

To determine the groups of the ATP molecule responsible for this capacity, we titrated C71G-PFN1 with ADP (Fig. S4). However, only at 1:8 (C71G:ADP), HSQC peaks of the unfolded state became completely disappeared and interestingly, most HSQC peaks with ADP at 1:8 are superimposable to those with ATP at 1:2. This indicates that ADP still has the capacity in shifting the equilibrium but has a weaker capacity than ATP. We also titrated with AMP (Fig. S5), but even with the concentrations up to 20 mM (1:400), AMP was still unable to completely convert the unfolded state. The results strongly suggest that the capacity of ATP in enhancing protein folding is dependent of triphosphate chain. Indeed, we titrated with adenosine and even with the highest concentrations of 5 mM due to its low solubility, no significant change was detected (Fig. S6).

By contrast, when titrated with triphosphate (PPP), at 1:1 (C71G:PPP), the HSQC peak intensity of the unfolded state also became significantly reduced (Fig. 2A and S7), while the 1D peak intensity of the methyl group of the unfolded state also became reduced while those of the folded state slightly increased (Fig 2B). At 1:2, HSQC peaks of the unfolded state became completely disappeared (Fig. 2C), and the intensity of the methyl group of the folded state further increased (Fig. 2B). Very strikingly, C71G-PFN1 with PPP at 1:2 has both HSQC (Fig. 2D) and 1D (Fig. 2B) spectra very similar to those with ATP at 1:2. However, further addition of PPP lead to the visible precipitation and NMR signals become too weak to be detectable. The results reveal that the capacity of ATP in enhancing folding mostly results from triphosphate but the isolated PPP also has a very strong ability to induce severe aggregation of C71G-PFN1 with significant packing defects due to the electrostatic screening effect (14). To assess whether the capacity in inducing folding results only from the electrostatic properties, we further titrated with sodium phosphate and chloride, and both failed to convert the unfolded state into the folded state with the concentration up to 5 mM for sodium phosphate (Fig. S8), and 10 mM for NaCl (Fig. S9), where NMR samples started to show visible precipitation. Noticeably, the addition of both salts appeared to induce the intensity reduction of both folded and unfolded peaks likely due to salt-induced aggregation (14). These results strongly imply that the inducing capacity of PPP is not just resulting from its high ionic strength because the ionic strength of sodium triphosphate is maximally 15-time stronger than sodium chloride while only 2.5-time stronger than sodium phosphate.

**Fig. 2.**
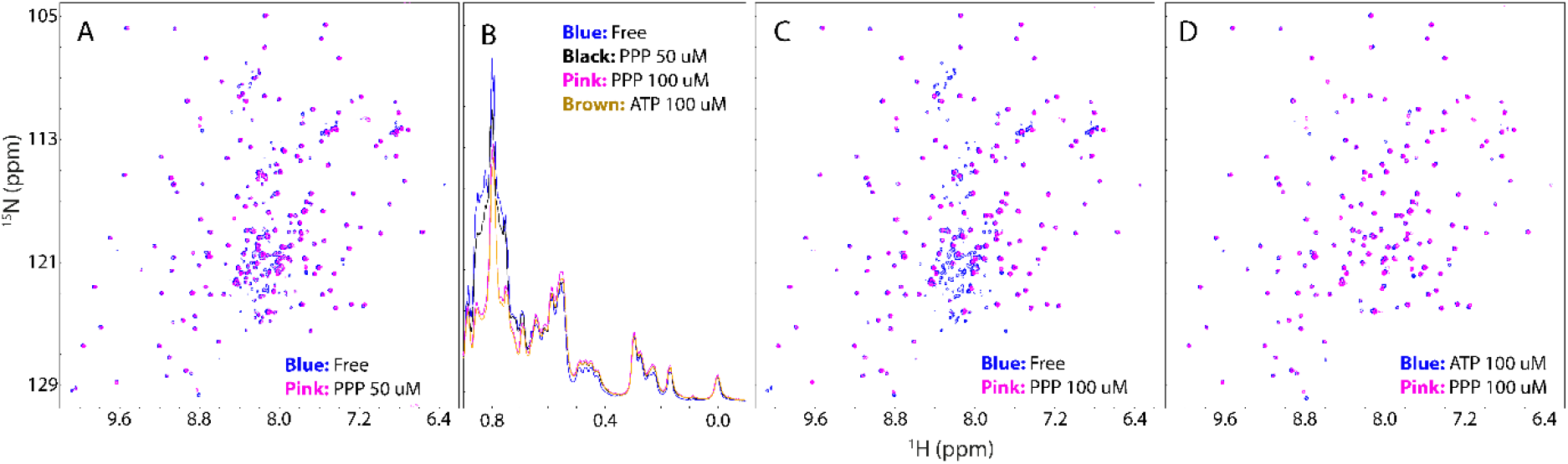
Triphosphate acid (PPP) shifts the conformational equilibrium to favor the folded state. Superimposition of HSQC spectra of ^15^N-labeled C71G-PFN1 at a concentration of 50 μM in the absence (blue) and in the presence of PPP (pink) at a molar ratio of 1:1 (A), and 1:2 (B). (C) Up-field 1D NMR spectra of C71G-PFN1 in the presence of PPP at different ratios. (D) Superimposition of HSQC spectra of ^15^N-labeled C71G-PFN1 in the presence of ATP at 1:2 (blue) or PPP (pink) at 1:2.

To assess whether the folding-inducing capacity of ATP and PPP is specific for C71G-PFN1, or a general phenomenon, we subsequently studied the nascent human superoxide dismutase 1 (hSOD1) associated with ALS (22,23). The 153-residue native hSOD1 with the zinc, copper and disulfide bridge Cys57-Cys146 adopts a β-barrel composed of eight antiparallel β-strands arranged in a Greek key motif (Fig. 3A), completely different from that of profilin-1 (Fig. 1A). However, the nascent hSOD1 lacking metal ions and disulfide bridge is highly unfolded and prone to aggregation in salted buffers as we previously characterized (23), which has HSQC (Fig. S10A) and 1D (Fig. S10B) spectra typical of an unfolded protein. Upon adding ATP to 1:2, no significant change was observed on HSQC (Fig. S10C) and 1D spectra (Fig. S10B). Nevertheless, when the ratio was increased to 1:4, well-dispersed HSQC peaks and very up-field 1D signals manifested, which was saturated at 1:8 (Fig. 3B, 3C and S11), indicating that the folded state was formed. Further addition of ATP to 1:4000 (20 mM) only induced some slight shifts of HSQC peaks (Fig S10D), indicating that unlike C71G-PFN1, ATP could not completely convert the unfolded state of hSOD1 into the folded state. Interestingly, many HSQC peaks of the folded states induced by ATP and by zinc are superimposable (Fig. 3D), but 1D signals have some differences (Fig. 3C), implying that they have similar backbone conformations but some difference in sidechain packing (24).

**Fig. 3.**
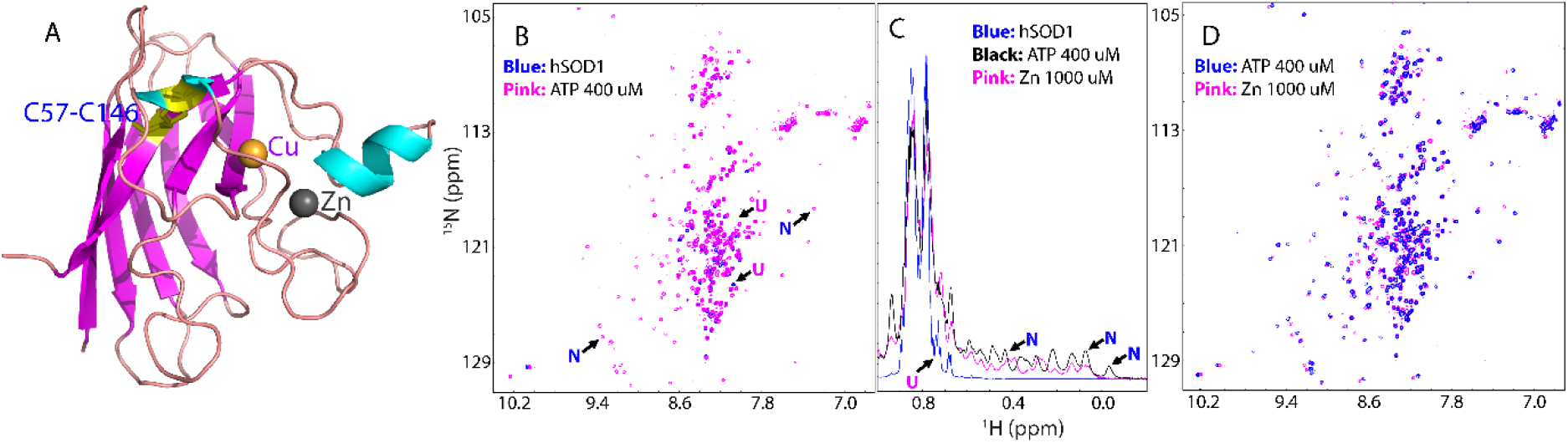
ATP can induce the folding of the unfolded nascent hSOD1. (A) The three-dimensional structure of the human superoxide dismutase 1 (PDB ID of 1MFM) with the cofactors zinc and copper cations displayed in spheres and disulfide bridge Cys57-Cys146 in sticks. (B) Superimposition of HSQC spectra of ^15^N-labeled hSOD1 at a concentration of 50 μM in the absence (blue) and in the presence of ATP (pink) at a molar ratio of 1:8. (C) Up-field 1D NMR spectra of hSOD1 at 50 μM in the absence (blue) and in the presence of ATP at 1:8 (pink) and zinc cation at 1:20 (black). (D) Superimposition of HSQC spectra of ^15^N-labeled hSOD1 in the presence of ATP at 1:8 (blue) and in the presence of zinc cation (pink) at 1:20.

We also found that like ATP, PPP could induce the folding of hSOD1. At 1:2, no significant change was observed on HSQC (Fig. S12A) and 1D (Fig. S12B) spectra. However, at 1:4, the folded population was formed as evidenced by the manifestation of the well-dispersed HSQC and up-field 1D peaks and this was mostly saturated at 1:8 (Fig. S12B, S12C and S13). Very interestingly, both HSQC (Fig. S12D) and 1D (Fig. S12B) spectra with PPP at 1:8 are very similar to those with ATP at 1:8, indicating the high similarity of the conformations induced by ATP and by PPP. Nevertheless. Also as observed on C71G-PFN1, further addition of PPP also triggered severe aggregation.

Previously we showed that the unfolded nascent hSOD1 could only be significantly induced to fold by zinc and at 1:20, this induction was saturated. However, even at 1:20, the unfolded state still co-existed with the folded state, and previously we obtained their residue-specific NMR conformations and dynamics (23). We also screened 11 other cations including Co^2+^, Ni^2+^, Cd^2+^, and Mn^2+^ commonly used to replace zinc in the native SOD1 for various structural studies, but all failed to significantly induce the folding of hSOD1 even at 1:40 (Fig. S14) where the samples started to precipitate. Very unexpectedly, ATP and PPP not only can induce folding of hSOD1 which was previously achieved only by zinc, but also have higher capacity than zinc. Therefore, we set to examine whether ATP and zinc together can completely shift the unfolded state. We added ATP to hSOD1 with the pre-existence of zinc at 1:20 (Fig. S15A and 15B). However, even with additional addition of ATP at 1:8, no significant change was observed for both HSQC (Fig. S15C and S15D) and 1D spectra (Fig. S15B).

So why can ATP completely convert the unfolded state of C71G-PFN1 but failed for hSOD1? Previously, we have collected HSQC-NOESY spectrum of the hSOD1 sample with zinc at 1:20, and found that no NOE cross peaks manifested for the two states, indicating that for hSOD1 with zinc cation, the exchange of the two states is much slower than the NMR time scale (23). In other words, the energy barrier separating the folded and unfolded states of hSOD1 are very large. Indeed, it was shown that the complete formation of the native hSOD1 needs further copper-load and covalent reaction to form disulfide bridge that can only be catalyzed by human copper chaperone for SOD1 (hCCS) (23).

We thus collected HSQC-NOESY spectrum for C71G-PFN1, and identified the cross peaks for the exchange of two states (Fig. S16). Therefore, by using the well-established NMR methods (25,26), we successfully mapped out the populations of 55.2% and 44.8% respectively for the folded and unfolded states, and the exchange rate of ~11.7 Hz (~85.5 milli-second) (Table S1). Moreover, we also determined the NMR dynamics on the ps-ns time scale by acquiring ^15^N backbone relaxation data T1, T2 and hNOE for both WT-PFN1 and C71G-PFN1 (Fig. S17), and subsequently performed the “model-free” analysis, which generates squared generalized order parameters, S^2^, reflecting the conformational rigidity on ps-ns time scale. S^2^ values range from 0 for high internal motion to 1 for completely restricted motion in a molecular reference frame (27,28). As shown in Fig 4A, the majority of the WT-PFN1 residues has S2 > 0.76 (with the average value of 0.89), suggesting that WT-PFN1 has very high conformational rigidity (Fig. 4B). By contrast, many residues of the folded state of C71G-PFN1 have S2 < 0.76 (Fig. 4A) (with the average value of 0.73) (Fig. 4C), revealing that even the folded state of C71G-PFN1 becomes more dynamic than WT-PFN1 on ps-ns time scale. Furthermore, the overall rotational correlation time (τc) of WT-PFN1 was determined to be 7.5 ns while that of C71G-PFN1 was 7.8 ns, implying that C71G-PFN1 becomes less compact, completely consistent with the difference of their translational diffusion coefficients (~1.12 ± 0.03 × 10^−10^ m^2^/s for WT-PFN1, and ~1.03 ± 0.02 × 10^−10^ m^2^/s for C71G-PFN1). The results imply that ATP induces protein folding by enhancing the intrinsic folding capacity of protein sequences, and consequently it failed to induce the complete conversion of the unfolded population of hSOD1 because this conversion needs the covalent formation of the disulfide bridge (23,24).

**Fig. 4.**
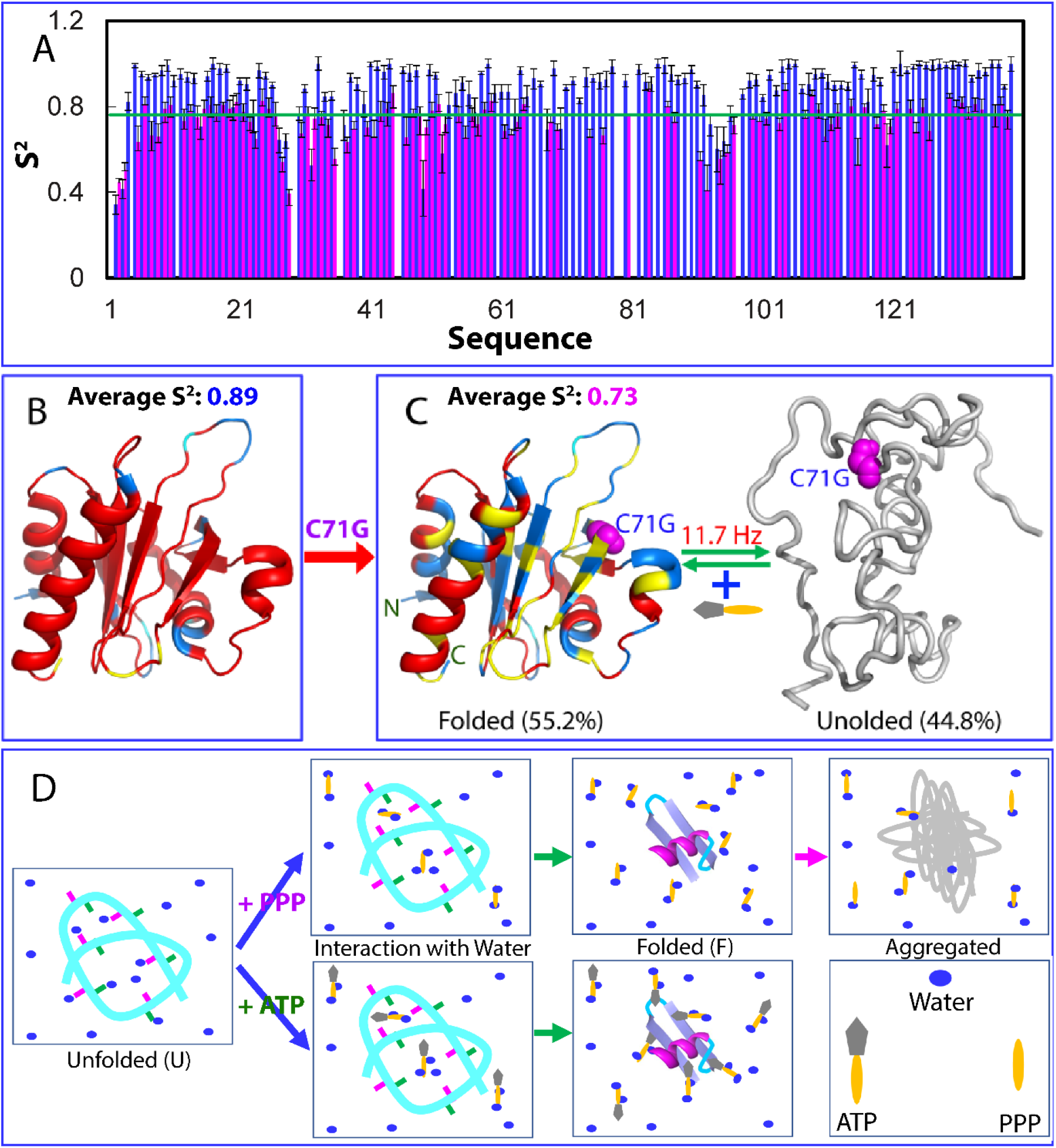
The mechanisms for ATP and PPP to modulate protein folding and dynamics. (A) Generalized squared order parameter (S^2^) of WT-PFN1 (blue) and the folded state of C71G-PFN1 (pink). The green line has a S^2^ value of 0.76 (average – STD). (B) The structure of PFN1 (PDB ID of 2PAV) with S^2^ values (with the average value of 0.89) of WT-PFN1 mapped on. (C) A diagram to show the co-existence of the folded (with the average S^2^ value of 0.73) and unfolded states of C71G-PFN1. S^2^ values are also mapped onto the folded state of C71G. Cyan is used for indicating Pro residues, and the yellow for residues with missing or overlapping HSQC peaks. Red is for residues with S^2^ values > 0.76 and blue for residues with S^2^ values < 0.76. The folded state with a population of 55.2% and unfolded state with a population of 44.8% have been experimentally mapped out to undergo a conformational exchange at 11.7 Hz. (D) A speculative model for protein folding induced by triphosphate (PPP) and ATP. For the unfolded state (U) of a protein, many backbone atoms are hydrogen-bonded with water molecules. Both triphosphate (PPP) and ATP can use their triphosphate chain to effectively attract water molecules out from hydrogen-bonding with the backbone atoms, thus favoring the formation of the folded state (F). However, due to the electrostatic screening effects produced by highly charged triphosphate, a protein exemplified by C71G-PFN1 with significant exposure of hydrophobic patches due to the defects in tertiary packing thus will become severely aggregated. By contrast, ATP can use its hydrophobic aromatic ring to dynamically interact with the exposed hydrophobic patches, which consequently not only functions to prevent from aggregation, but also to increase the thermodynamic stability.

We also measured the thermodynamic stability by differential scanning fluorimetric (DSF) method (18). Interestingly, WT-PFN1 has a melting temperature (Tm) of ~56 degree while the addition of ATP even up to 20 mM triggered no significant change (Fig. S18A), consistent with our previous NMR results that ATP has no significant interaction with WT-PFN1 (18). By contrast, C71G-PFN1 has no cooperative unfolding signal (Fig S18B), likely due to the absence of tight tertiary packing or/and co-existence of two states. Interestingly in the presence of ATP of 20 μM (a molar ratio of 1:2), a cooperative unfolding signal was observed with Tm of 32 degree. Addition of ATP to 1 mM (1:100) led to the increase of Tm to 38 degree, and further increase to 40 degree at 20 mM (Fig. S18C). By contrast, we performed many DSF measurements of C71G with PPP at various ratios and obtained unfolding curves with no cooperative unfolding signals but high noises, implying that although ATP and PPP have similar capacity in inducing folding, PPP failed to enhance tertiary packing of the folded state or/and triggered dynamic aggregation even before the visible precipitation. Indeed, previously we found that a protein still had native-like NMR spectra although its tight packing was disrupted to different degrees (24).

In this study, for the first time we discovered that ATP has a general capacity in inducing protein folding at very low ATP/protein ratios which can be satisfied in most cells. Previously, the best-known molecule with the general capacity is the natural osmolyte, trimethylamine N-oxide (TMAO) (4,29). However, to significantly induce the folding of the reduced RNase T1 at 10 μg/ml, TMAO concentrations needed to reach ~1.2 M (29). Therefore, ATP has the strongest efficiency known so far in inducing protein folding. It is also of fundamental significance to identify that the inducing capacity is from triphosphate which was previously proposed to act as a key intermediate for various prebiotic chemical reactions, which allowed the Origin of Life (30). Our current study thus decodes a previously-unknow rationale why nature chose phosphate as the building element for the Origin of Life.

So what could be the mechanism for PPP to induce protein folding so effectively? Recently, it was proposed that the hydrogen bonding and solvation of the backbone plays a key role in protein folding (3–5). The unfolded state (U) will be favored if the backbone atoms are highly hydrogen-bonded with water molecules, while the native state (N) will be favored if the backbone atoms are involved in forming intramolecular hydrogen-bonds (4). Here as illustrated by Fig. 4D, we propose that like TAMO, both triphosphate and ATP dynamically interacts with the protein surface, and very efficiently attracts water molecules out from hydrogen-bonding with the backbone atoms. Consequently, the backbone atoms will be shifted to form the intramolecular hydrogen bonds to favor protein folding. Indeed, previously we found that triphosphate has unusually high capacity in interacting with water molecules (17). However, because of being highly charged, triphosphate at high concentrations will unavoidably provoke significant screening effect to trigger aggregation for the unfolded or partially-unfolded states of proteins as exemplified by C71G-PFN1 and nascent hSOD1 with significant exposure of hydrophobic patches (14). This may also explain why in modern cells, the concentrations of triphosphate are extremely low. Remarkably, however, nature has successfully solved this problem by joining adenosine and triphosphate together to form ATP, which not only retains the capacity of PPP in efficiently inducing protein folding, but also eliminates the capacity in severely triggering aggregation by using the aromatic ring of adenosine to dynamically interact with the exposed hydrophobic patches. Strikingly ATP further acquired a novel ability to increase the thermodynamic stability of the proteins with the defects in tertiary packing such as C71G-PFN1 This observation thus offers a principle to engineer three key functions into one small molecule to treat aggregation-associated ageing and diseases, as well as for other applications.

Now with our current results, ATP has been established to own the ability to modulate both sides of protein homeostasis: inducing folding and inhibiting/dissolving aggregation with various energy-independent mechanisms (12,13,17–20), which might play a central role in the emergence of cells during prebiotic evolution. In the model cells, ATP further acts to drive the energy-dependent chaperone and disaggregase machineries to control protein homeostasis. Therefore, in addition to controlling protein homeostasis associated with ageing and diseases in modern cells, ATP appears also to play the irreplaceable key roles in the Origin of Life.

## Acknowledgments

This study is supported by Ministry of Education of Singapore MOE Tier 1 R-154-000-B45-114 Grant and Tier 2 MOE2015-T2-1-111 (to J.S.).

